# A real-time cellular thermal shift assay (RT-CETSA) to monitor target engagement

**DOI:** 10.1101/2022.01.24.477382

**Authors:** Tino W. Sanchez, Michael H. Ronzetti, Ashley E. Owens, Maria Antony, Ty Voss, Eric Wallgren, Daniel Talley, Krishna Balakrishnan, Ganesha Rai, Juan J. Marugan, Sam Michael, Bolormaa Baljinnyam, Noel Southall, Anton Simeonov, Mark J. Henderson

## Abstract

Determining a molecule’s mechanism of action is paramount during chemical probe development and drug discovery. The cellular thermal shift assay (CETSA) is a valuable tool to confirm target engagement in cells for a small molecule that demonstrates a pharmacological effect. CETSA directly detects biophysical interactions between ligands and protein targets, which can alter a protein’s unfolding and aggregation properties in response to thermal challenge. In traditional CETSA experiments, each temperature requires an individual sample, which restricts throughput and requires substantial optimization. To capture the full aggregation profile of a protein from a single sample, we developed a prototype real-time CETSA (RT-CETSA) platform by coupling a real-time PCR instrument with a CCD camera to detect luminescence. A thermally stable Nanoluciferase variant (ThermLuc) was bioengineered that withstood unfolding at temperatures greater than 90 degrees Celsius and was compatible with monitoring target engagement events when fused to diverse targets. Utilizing well-characterized inhibitors of lactate dehydrogenase alpha, RT-CETSA showed significant correlation with enzymatic, biophysical, and other cell-based assays. A data analysis pipeline was developed to enhance the sensitivity of RT-CETSA to detect on-target binding. The RT-CETSA technology advances capabilities of the CETSA method and facilitates the identification of ligand-target engagement in cells, a critical step in assessing the mechanism of action of a small molecule.

**Significance:** Validating target engagement is a critical step when characterizing a small molecule modulator. The cellular thermal shift assay (CETSA) is a common approach to examine target engagement, as alterations in the thermal stability of a protein can be conferred by ligand binding. An advantage of CETSA is that it does not require modification of the protein target or small molecule. Major limitations are the throughput and ease-of-use, as the traditional detection method uses western blots, which limits the number of samples that can be processed. Higher-throughput CETSA methods have been developed but are performed at a single temperature and require target-specific optimization. We developed a high-throughput real-time CETSA to circumvent these challenges, providing a rapid and cost-effective strategy to assess on-target activity of a small molecule in living cells.

## Introduction

The cell is a complex environment with numerous, tightly controlled biochemical reactions and interactions between cellular components (1). Proteins are central to many cellular processes, and there is indispensable value in identifying small molecules that target the proteome with high specificity and affinity Confirming the engagement between a protein target and small molecule under physiologically relevant conditions poses a substantial challenge in early-stage drug discovery and probe development. Most strategies to study target engagement are labor intensive, low-throughput, or do not provide evidence of ligand-target binding in a physiological, cellular environment (2). The gold standard for target engagement remains co-crystallization of target and ligand using x-ray crystallography, but this methodology remains highly complex, is not amenable to all target classes, and is not suitable for testing large numbers of compounds. Sensor-based biophysical methods like isothermal calorimetry and surface plasmon resonance (SPR) detect direct target binding but implement simplified acellular conditions and require significant amounts of purified protein and optimization (3, 4). Thermal-shift based biochemical methods such as differential scanning fluorimetry (DSF) also detect ligand-induced thermal shifts using recombinant protein with hydrophobic dyes or intrinsic protein fluorescence (nanoDSF) (5–7). None of these approaches account for complexities found in cells, including membrane barriers and the potential for off-target binding.

The cellular thermal shift assay (CETSA) allows for the study of target engagement with a small molecule or biomolecule in intact cellular environments, linking observed phenotypic responses with a compound’s molecular target (8, 9). Thermal shift assays rely on detecting thermodynamic stabilization of a protein resulting from ligand binding adding discrete bond energy and shifting the Gibbs free energy of the system. This shift in system energy can be detected by measuring the aggregation properties of the target protein when a thermal challenge is applied (10). Traditionally, the CETSA method is performed as a lytic endpoint assay, requiring individual samples to be prepared for each temperature or compound concentration. After incubating cells with a compound of interest, samples are heated to discrete temperatures and the unfolded aggregated protein and cellular debris is removed by centrifugation. Remaining soluble protein is then measured using a protein detection method, most commonly western blot, though more recent methods have utilized mass spectrometry (11–13).

Modernizing the CETSA approach for higher-throughput drug discovery applications requires the development of alternative CETSA-compatible detection methods, such as fluorescent and bioluminescent reporters (14–16). One recent example was the development of a homogenous bioluminescent assay using a split Nano luciferase reporter (SplitLuc CETSA). By appending a small HiBiT-based peptide reporter tag to the target and adding the complementary LgBiT protein with a furimazine substrate, SplitLuc CETSA allows for measurements in 384- and 1536-well microplates, providing an improvement in throughput (17). In a similar approach, native Nano luciferase (NLuc) was fused to different targets to measure ligand-induced thermal shifts by detecting changes in the luminescent signal (NaLTSA) (18). To date, all CETSA methods perform end-point measurements, so independent samples are needed at each discrete temperature being assessed (19). Moreover, classical CETSA analysis relies on single parameter methods using a sigmoidal fit of thermal response curves to either calculate the midpoint aggregation temperature (T_agg_) or area-under-curve (AUC). Indeed, the limitations of high-throughput CETSA methods like SplitLuc and NaLTSA (NLuc CETSA) and the applied methodology to analyze thermal profiles, stem from the requirement to choose whether to examine either the dose-response of small molecules at a single temperature, or a single concentration of drug over a range of temperatures (17–22).

We reasoned that a real-time CETSA (RT-CETSA) assay that captures the full thermal melt profile within living cells would bridge the CETSA information gap and enable the acquisition of high-throughput information-rich data across a temperature ramp from a single sample. We further reasoned that using NLuc would be a favorable reporter tag for target proteins due to the low background luminescence, bright signal and no interference with intrinsic fluorescence of small molecules (23). However, NLuc has an aggregation temperature (T_agg_), that ranges between 55 to 60 °C, precluding its use as a reporter for a portion of the proteome (23–25). We hypothesized that a thermally stable NLuc variant would reduce the reporter’s propensity to drive aggregation due to thermal unfolding, and that the NLuc variant LgBiT (11S), which was engineered to exhibit improved intracellular stability (23), might exhibit the increased thermal stability required. Herein we describe the bioengineering of thermally stable luciferase variants (ThermLuc) and the creation of a proof-of-concept RT-CETSA detection device. To quantify thermal unfolding in RT-CETSA, we also developed a novel approach using baseline-corrected thermal unfolding curves from MoltenProt, a recently developed analysis pipeline that produces nonlinear fits of protein unfolding, and goodness of fit tests between two models to determine thermal stabilizing molecules (26). Herein, the RT-CETSA technology platform is described and validated using lactate dehydrogenase alpha (LDHA)-ThermLuc fusions and a diverse set of pyrazole-based LDHA inhibitors (27).

## Results

### Bioengineering Thermally Stable Luciferase Fusion Proteins for Improved Detection of Ligand-Induced Thermal Stabilization

A critical component of the proposed RT-CETSA approach is the creation of a thermally stable luminescent reporter that continuously produces signal throughout a CETSA temperature ramp but does not drive reporter-led aggregation due to its own thermal unfolding. We hypothesized that an engineered NLuc using a LgBiT-based protein would be more thermally stable than traditional NLuc. First, we measured the stability of purified LgBiT and HiBiT fragments against a thermal challenge using nanoDSF. Thermal shift experiments revealed that combining LgBiT and HiBiT resulted in a significant T_agg_ shift to 73.8 °C compared to 45.2 °C for NLuc (using the corresponding 156 and NP fragments) (Fig. 1A). This provided the rationale to generate a plasmid encoding LgBiT and HiBiT assembled into a single fusion protein for heterologous expression in cells. Six constructs containing varying lengths of Gly-Ser linkers between LgBiT and HiBiT were transfected into HEK293T cells and assessed for both luminescence (Fig. 1B) and thermal stability (Fig. 1C). Although total luminescence was lower for samples expressing the LgBiT/HiBiT fusion proteins, they exhibited greater thermal stability, with an increase in T_agg_ from 63 °C (NLuc) to >90 °C. The construct with a single Gly-Ser peptide linker between luciferase fragments improved thermal stability compared to NLuc but to a lesser degree than the longer Gly-Ser linkers. We selected the fusion protein containing six Gly-Ser repeats, hereafter referred to as ThermLuc, for subsequent experiments measuring luminescence and thermal stability (Supplementary Table 1).

**Fig. 1:**
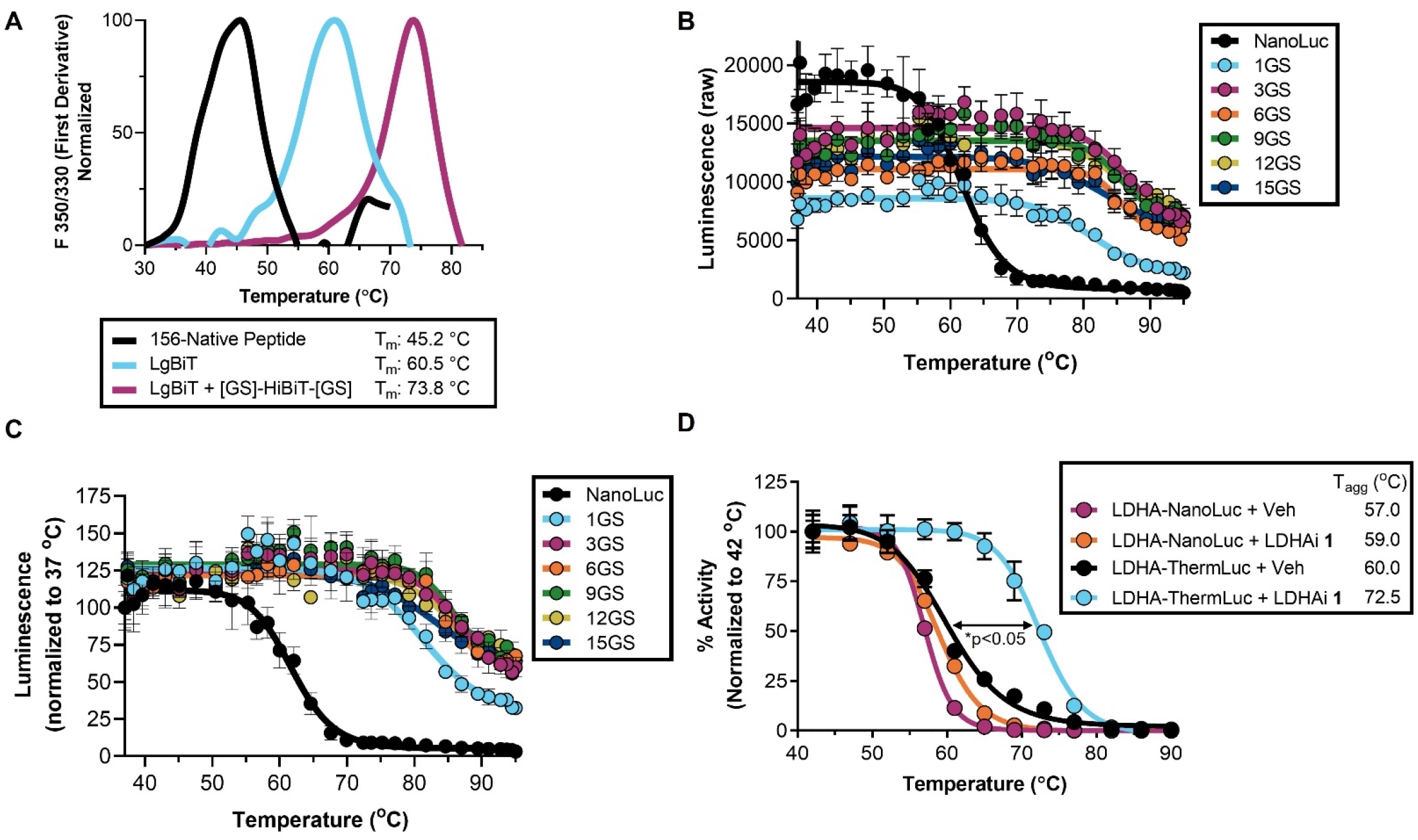
Engineering a thermally stable NLuc variant (ThermLuc). (**A**) Nano-differential scanning fluorimetry (nanoDSF) analysis of the thermal stability of reconstituted traditional NLuc (split into 156 and native peptide fragments), LgBiT, and LgBiT plus GS-HiBiT-GS peptide. (**B, C**) LgBiT and GS-HiBiT-GS fragments were combined into a single fusion protein and transiently expressed in HEK293-T cells. Different Gly-Ser linker lengths to connect the fragments were examined. Graphs represent (**B**) raw luminescence values (mean ± SD, n=4) and (**C**) luminescence normalized to the 37 °C signal (mean ± SD, n=4). (**D**) The thermal shift expected from the binding of an LDHA inhibitor (LDHA_i_) **1** is masked when using a LDHA-NLuc fusion, but detectable with a LDHA-ThermLuc fusion (mean ± SD, n=4).

To assess the utility of ThermLuc as a CETSA reporter, we created a fusion with LDHA, a 35 kDa soluble protein that aggregates in the low 60’s °C. Small molecule-induced stabilization with 10 μM of LDHA inhibitor (LDHA_i_) **1** was nearly undetectable under NaLTSA conditions, which is consistent with the unfolding of NLuc driving aggregation and masking the thermal stabilization effect of target engagement (Fig. 1D). Swapping NLuc with the ThermLuc reporter slightly increased the apparent T_agg_ by 3.0°C but showed a significant increase in thermal stability of the LDHA fusion with compound **1** (ΔT_agg_ = 12.5 °C). This data supports the hypothesis that improved thermal stability of the luminescent reporter can unmask ligand-induced thermal stabilization of the target-of-interest, as the reporter no longer drives aggregation of the fusion protein due to its own thermal unfolding.

### Real-Time CETSA Overview

To explore whether the entire aggregation profile of a target protein within its natural cellular environment could be monitored during heating, we pursued an RT-CETSA procedure utilizing the bioengineered ThermLuc protein (Fig. 2). The method proceeds with the following steps: 1) cells are transfected with a plasmid encoding the target of interest (TOI) fused to ThermLuc, 2) cells expressing the ThermLuc fusion protein are dispensed into PCR plates and ligands are added, and 3) the luciferase substrate furimazine is added and luminescence is recorded kinetically as temperature is increased stepwise (e.g. 1 °C increments from 37 °C to 90 °C) to define the melt profile. The RT-CETSA method inherently requires a detection device that couples precise temperature control with a sensitive luminescence detection system (Supplementary Fig. S1). Modern qPCR machines are well suited for temperature control as they use thermal blocks with sub-centigrade precision and uniform heating across samples, but these machines are exclusively paired with detection systems optimized for fluorescence quantitation; there are no options available for luminescence detection. Therefore, we adapted a high-throughput qPCR instrument (LightCycler 480 II, Roche), removed filters from the light path, and swapped the stock fluorescence camera with an Orca R2 CCD to measure ThermLuc output signal while heating (Supplementary Fig. S1). Next, an assessment of the thermal unfolding profiles of ThermLuc-targets began with automated pixel intensity analysis in MATLAB to extract raw luminescence values, followed by analysis of the melting curves using T_agg_, AUC, and a novel nonparametric thermal curve analysis method to determine ligand-induced stabilization in RT-CETSA.

**Fig. 2:**
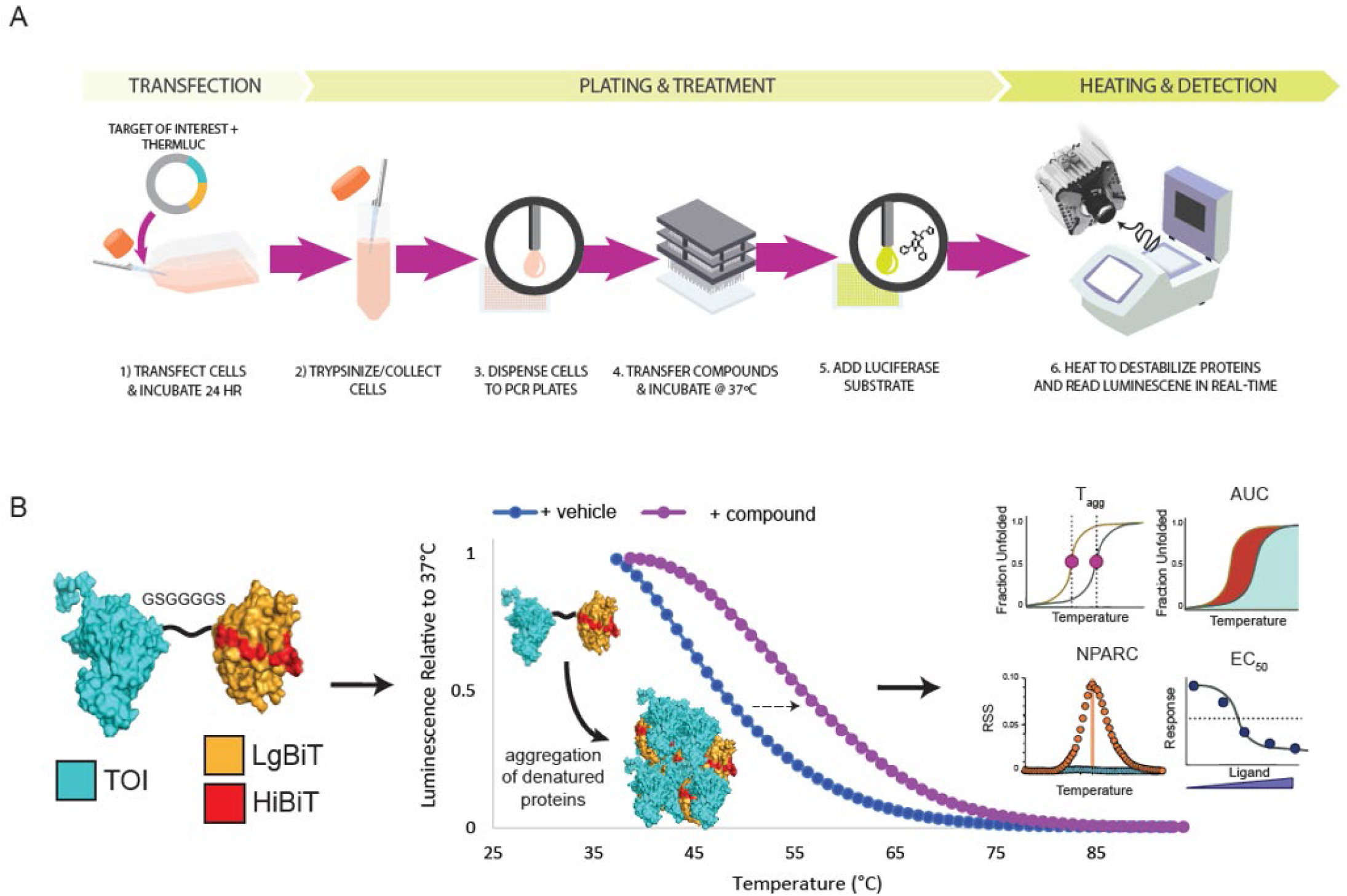
Real-time CETSA (RT-CETSA). (**A**) Schematic overview of the RT-CETSA approach. (**B**) Depiction of aggregation of ThermLuc-fusion protein at elevated temperature, resulting in loss of luminescence. The entire melt profile can be measured from a single sample and target engagement can be detected as a change in apparent T_agg_, AUC and novel nonparametric curve analyses (NPARC) further described herein.

### Real-Time Monitoring of Heat-Induced Aggregation of NLuc Variant Fusions

The melting behavior of NLuc and ThermLuc luciferase fusion constructs were compared using a real-time detection approach. ThermLuc fusion proteins with three to fifteen Gly-Ser linker repeats between the HiBiT and LgBiT fragments showed significantly higher aggregation temperature (T_agg_) compared to native NLuc when captured in real-time (Supplementary Fig. S2A). Moreover, the reporter with a single Gly-Ser linker had a T_agg_ in between native NLuc and the ThermLuc constructs with longer linkers (Supplementary Fig. S2A), consistent with behavior in the endpoint luminescence lytic CETSA experiment (Fig. 1C). Notably, the apparent T_agg_ in the RT-CETSA system may not align with T_agg_ values calculated using traditional luminescence or immunoblotting detection. Traditional CETSA utilizes a 3-min hold at a single temperature to calculate T_agg_, as opposed to a rapid ramping and recording over a temperature range as in the real-time protocol. Additionally, the apparent T_agg_ in RT-CETSA is impacted by extrinsic factors that also contribute to luminescent signal, such as heat-induced substrate decomposition, which occurs at temperatures greater than 60 °C (Supplementary Fig. S2B). The decay rate of luminescence is largely attributable to temperature effects; maintaining 37 °C throughout the RT-CETSA experiment shows minimal signal loss (Supplementary Fig. S2C). Changing experimental conditions, such as increasing the hold time at each temperature, alters the apparent melting behavior (Supplementary Fig. S2D). RT-CETSA melt profiles therefore are not expected to align with absolute T_agg_ values calculated using traditional CETSA; rather, the primary goal of RT-CETSA is to identify thermal shifts due to ligand-induced stabilization.

The nature of real-time measurements using a luciferase reporter requires the presence of a reporter substrate (furimazine) from the beginning timepoint, which presents additional experimental design considerations. The commercially available furimazine substrate typically used in NaLTSA experiments (Promega NanoGlo Substrate) is formulated in undisclosed chemical matter, so we examined whether using an alternative common solvent such as DMSO would be compatible with RT-CETSA. The melting profile of the ThermLuc fusion proteins and NLuc was very similar to that observed with the commercially available furimazine (Supplementary Fig. S2A, C). An additional consideration of including furimazine during heating is the potential for negative effects from the substrate on cell physiology. We performed a viability assay and found that cell growth was significantly impaired with a 48-hour treatment of the commercially available furimazine when used at concentrations 0.5X and higher, but not the DMSO formulated furimazine up to 50 μM (Supplementary Fig. S2E). To further explore effects of furimazine on live cells, we performed a cell health screen using 0.005 – 100 μM of the DMSO formulated furimazine in a SYSTEMETRIC cell health screening platform (AsedaSciences). The overall score in this assay placed furimazine in the ‘low cell stress’ category, but effects on reactive oxygen species, membrane permeability and nuclear membrane permeability were detected (Supplementary Fig. S2F). Importantly, the presence of furimazine during the heating step did not significantly alter target engagement and thermal stabilization of LDHA as conferred by an LDHA inhibitor (Supplementary Fig. S2G).

We next considered the rate of temperature increase that would bring the RT-CETSA system towards thermal equilibrium and properly capture aggregation profiles. Traditional CETSA protocols heat samples for 3 – 3.5 mins, but there are a limited number of experiments that address whether this long incubation period is required. We performed endpoint lytic CETSA with LDHA-ThermLuc and found that the melting profiles were similar after 30 sec and 3.5 min of heating (Supplementary Fig. S2H). Next, we examined the rate of unfolding using RT-CETSA. Applying 72 °C of heat brought about thermal unfolding of LDHA rapidly, with most of the aggregation occurring within 30 sec (Supplementary Fig. S2I). These results suggest that long incubation times, like those described in the original CETSA protocols, are not required to detect compound-induced thermal shifts for all targets.

### Target Engagement of LDHA-ThermLuc in RT-CETSA

The ability to detect small-molecule induced thermal shifts using the RT-CETSA method was first examined using an LDHA-ThermLuc fusion and a well-characterized inhibitor (compound **1**). Tangential work on thermal proteome profiling, a version of CETSA relying on mass-spectrometry as the method of readout, has used nonparametric analysis of response curves (NPARC) to integrate goodness of fits of the entire thermal response curve as an alternative to summary statistics like T_agg_. NPARC is more sensitive and specific to ligand-induced thermal stabilization than the melting point and other single-parameter values, and so we sought to integrate this method for analyzing RT-CETSA data. Using MoltenProt analysis of the unfolded protein, we measured ligand-induced stabilization of LDHA across a temperature gradient (Fig. 3A). To assess reproducibility of the RT-CETSA method, we examined 192 replicate wells of either DMSO or LDHA_i_ **1** scattered across a 384-well plate. We observed a range of the fraction-unfolded values across the plate (DMSO: 0.063 ± 0.006 [9.45% CV], LDHA_i_ **63**: 0.051 ± 0.005 [9.51% CV]) which may be attributable to experimental variability in cell number across wells (Fig. 3B). Despite this variability, the starting luminescence did not affect the percent melt of LDHA-ThermLuc, and LDHAi **1** stabilization remained consistent, highlighting an advantage to capturing kinetic RT-CETSA data for every sample, where each sample can be normalized to its starting signal before heating (Fig. 3B). Next, we compared each analysis method applicable to RT-CETSA data (T_agg_, AUC, NPARC) using the Z’ assay reproducibility statistic. Using DMSO vehicle as a negative control (N=192) compared to LDHAi **1** treated wells as a positive control (N=192), we found the best performing Z’ to be NPARC [0.72] >> AUC [0.55] >> T_agg_ [0.52] (Fig. 3C). The RT-CETSA thermal unfolding curve fitting approach also had acceptable signal windows with these controls (T_agg_: 6.23, AUC: 7.57, NPARC: 20.40) and a repeatable Δ4 °C compound-induced thermal shift across all wells (Fig. 3C).

**Fig. 3:**
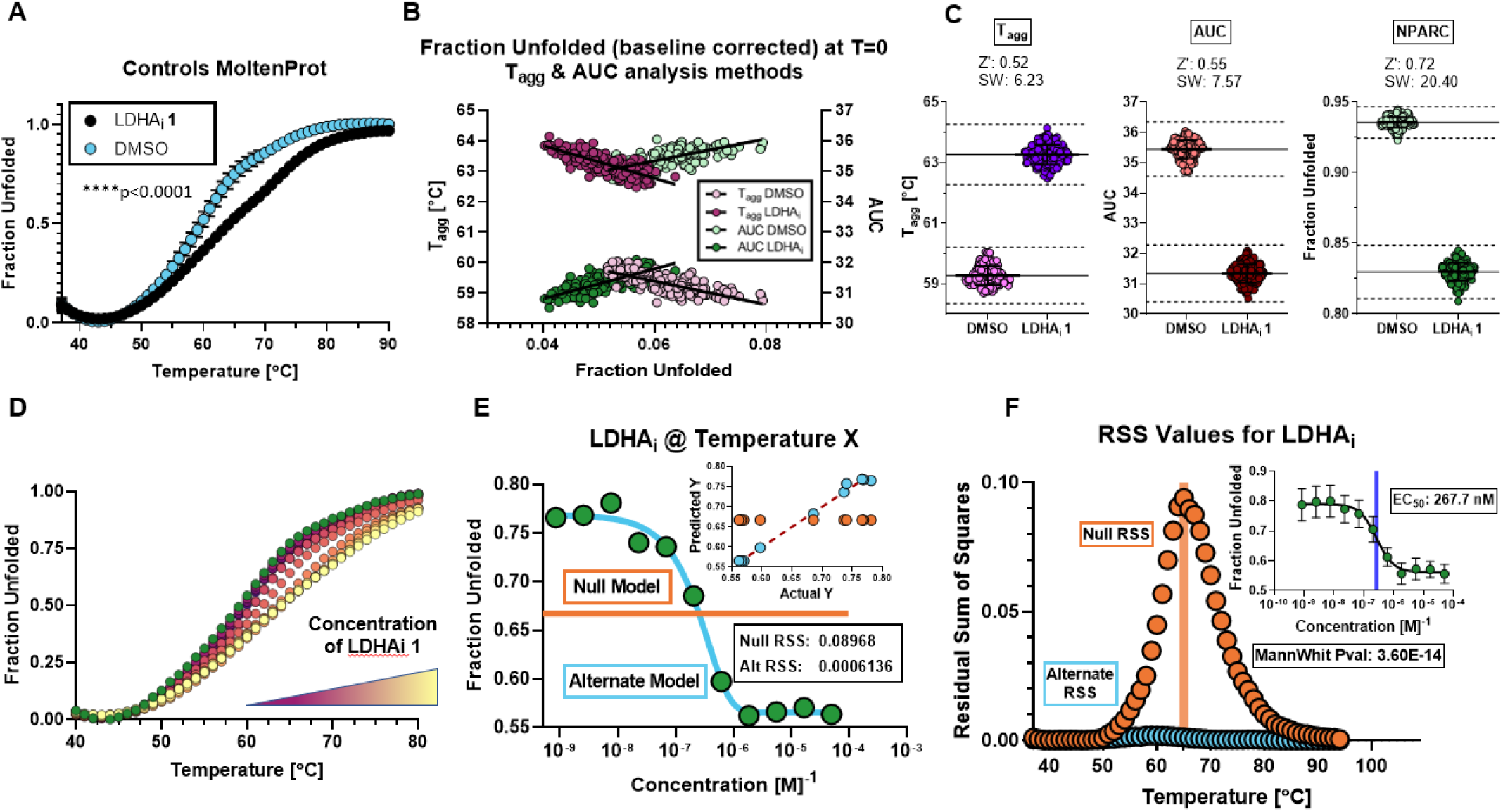
RT-CETSA thermal unfolding curves for LDHA-ThermLuc incubated with vehicle or LDHA_i_ **1** after processing with MoltenProt. (**A**) Data shown is 3 biological replicate plates with N=192 for each group per plate with standard deviation error bars. (**B**) Distribution of vehicle and LDHA_i_ **1** baseline corrected fraction unfolded values at the first temperature point against T_agg_ and AUC parameters from 3 biological replicate plates with N=576 for each group. (**C**) Distributions of positive and negative controls using LDHA-ThermLuc are used to determine the Z’ statistic and signal window using T_agg_, AUC, and NPARC methods of analysis. Solid lines represent the means of each group, and dashed lines represent the ±3*SD for each control group. (**D**) Thermal dose-response curves of LDHA_i_ **1** used for processing with MoltenProt and RT-CETSA scripts from the LDHA_i_ experiment. (**E**) Goodness-of-fit tests for dose-response data at each discrete temperature point (green circles) are performed with a null (linear fit with slope constrained to 0, shown as orange) and alternative model (four-parameter log logistic fit), from which residual sum of squares (RSS) are calculated. Predicted vs. actual Y values for each model are shown in the graph inset, showing the high degree of fit with the alternate model (blue circles) and the higher residuals for the poorly fit linear model (orange circles) (**F**) RSS values for the null (orange) and alternate (blue) models are plotted and the difference in RSS between models is calculated. The point of maximal RSS difference (shown as an orange bar) is used to determine EC_50_ of the compound, shown as the blue bar in the graph inset of concentration-response values.

To analyze real-time thermal melt profiles of LDHA inhibitors, we modified previous NPARC methods to create goodness of fit models for dose-response thermal melting curves, like the LDHA_i_ **1** dose-response set in Fig. 3D, by fitting every concentration at every temperature with a null (linear fit with a slope of 0) and alternate (4-parameter log-logistic fit) model, and then calculating residual sum of squares (RSS) for each model fit (Fig. 3E). The null fit is the theoretical model for when there is no dose-dependent stabilization of a target, in other words no significant change to the melting profile of the target with ligand compared to a vehicle control. Thus, we expect to see a poorer fit of the null model, as compared to the alternate model, when dose-dependent stabilization of the target occurs, as illustrated by the actual vs predicted Y values for an LDHA_i_ at a temperature near the aggregation temperature for LDHA (Fig. 3E inset). A goodness of fit test of these residuals is performed with the nonparametric Mann Whitney U test against the model RSS values to detect significant binders (thermal shift/stabilization), and an EC_50_ is calculated for binders by analyzing dose-response curves at the point of maximal RSS difference using a 4-parameter log-logistic fit (Fig. 3F). We note that the point of maximal difference between null and alternate models is often at or near the T_agg_ for the target (Fig. 3F).

### RT-CETSA Target Engagement of LDHA Shows Compatibility Across Platforms

We benchmarked the RT-CETSA platform for its ability to guide structure-activity-relationship (SAR) studies on a set of 29 previously identified LDHA inhibitor analogs by comparing activity in complementary biochemical, biophysical, and phenotypic assays (Supplementary Table 2). Experiments were performed in dose response ranging from micromolar to sub-nanomolar concentrations to determine EC_50_ values (Supplementary Movie S1). The compounds were pre-incubated for 1 hour in RT-CETSA; notably, a time course examination of LDHAi **1** showed diminished target engagement when compound pre-incubation was reduced to 15 minutes (Supplementary Fig. S2J). In three independent experiments, 29 inhibitors were tested using the RT-CETSA method; EC_50_ values were calculated using T_agg_, AUC and the modified NPARC analysis that was developed for RT-CETSA. These three RT-CETSA analysis methods were further compared to SplitLuc CETSA (isothermal heating performed at 61, 65, and 69 °C) and other biophysical and biochemical assays (Fig. 4A, Supplementary Fig. S3A-C). For all LDHA inhibitors analyzed, the NPARC method derived a range of baseline-corrected fraction unfolded values from ~0.5 to ~0.8 (Fig. 4B). EC_50_ values were calculated using the point of maximal difference between null and alternate models. In addition to a superior Z’ median for assay reproducibility, NPARC analysis was more sensitive and specific to ligand-induced thermal stabilization than T_agg_ or AUC (Fig. 4A). For example, compound **19** had an EC_50_ value of 14.6 nM in RT-CETSA using NPARC but was inactive at 50 μM using conventional T_agg_ and AUC analysis. When comparing across different assays, most of the inhibitors with low nanomolar EC_50_ values in the biochemical assay using recombinant LDHA, also had nanomolar EC_50_ values in the cellular-based SplitLuc CETSA and RT-CETSA target engagement assays using NPARC analysis. For example, compound **18** had a biochemical EC_50_ of 63 nM with recombinant LDHA, a SplitLuc CETSA EC_50_ of 100 nM and an RT-CETSA EC_50_ of 14 nM using modified NPARC analysis. The rank order of the compounds as inhibitors of LDHA were all significantly correlated in the RT-CETSA assays using all three analysis methods when compared to SplitLuc and other biophysical/biochemical data (Fig. 4C). RT-CETSA however, showed more potent response profiles compared to the SplitLuc CETSA approach that uses a 3.5 min heating step and endpoint analysis (Supplementary Fig. S3D). All three methods for analysis of RT-CETSA were tested for their replicate-experiment minimum significant ratio (MSR), although only NPARC (MSR: 2.32) and AUC (MSR: 2.67) methods were shown to reach acceptable reproducibility (Fig. 4D, Supplementary Fig. S3E). We observed good correlation between potency of target engagement for RT-CETSA using NPARC and traditional CETSA using endogenous protein; however, two compounds showed more potent stabilization in RT-CETSA (Fig. 4E). Notably, these were compounds that exhibit shorter residence time, as measured by SPR, which may reflect an increased ability to detect engagement of compounds with faster off rates using the RT-CETSA method.

**Fig. 4:**
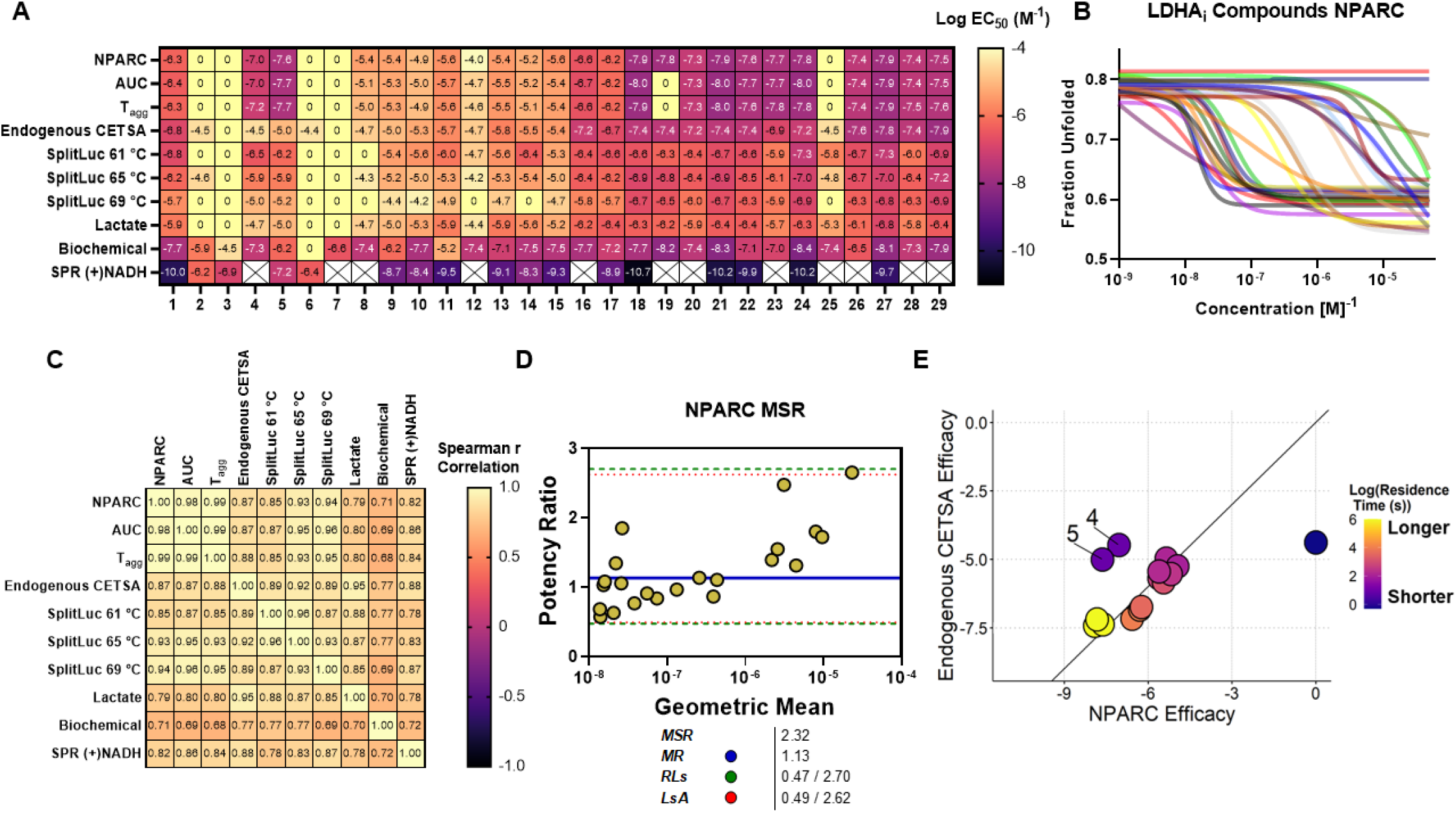
Correlative analysis of LDHA inhibitors. **A)** EC_50_ values (Log M) for plate containing 29 LDHA inhibitors (n=3 replicates) analyzed using the following methods: RT-CETSA, endogenous CETSA, SplitLuc CETSA, lactate assay, biochemical assay and SPR. Compounds with no detected binding are annotated as “0”, and compounds with no data for a particular method are annotated with a blank square. Data shown is the mean of 3 replicates of RT-CETSA experimental data. (**B**) Dose-response curves for all LDHA inhibitors when using nonparametric analysis of thermal dose-response curves. NPARC derives fraction unfolded values for EC_50_ calculation at the point of maximal difference between null and alternate models, presenting a range of fraction unfolded values from ~0.8 for low concentrations of stabilizing small molecule and ~0.5 for higher concentrations. (**C**) Spearman correlations of the compound rank order shows significant correlation among methods tested. All correlations were statistically significant (p<0.005, two-tail). (**D**) Testing of the minimum significant ratio (MSR) and related parameters further characterize the high reproducibility of potency estimates from the RT-CETSA method. The mean ratio (MR) is shown as a solid blue line, Limits of Agreement (LsA) in dashed red lines, and ratio limits (RL) in dashed green lines. (**E**) Examination of EC_50_ values for compounds when tested in RT-CETSA versus SplitLuc CETSA. The color of the points indicates residence time, as calculated by surface plasmon resonance (SPR).

### Expanding the Utility of RT-CETSA to Simultaneously Measure the Thermal Profile of Multiple Targets

The significant correlation and compatibility of RT-CETSA to identify ligand-induced stabilization and target engagement of a well-characterized set of LDHA_i_ analogs validates this method as a high-throughput screening platform for future SAR studies. We hypothesized that we could leverage the RT-CETSA methodology to monitor the real time melt profile of different targets simultaneously across a single plate. As a proof-of-concept, we used RT-CETSA to measure the thermal melt profile of nine different target-ThermLuc fusion proteins (Supplementary Fig. S4A). This set included immunotherapeutic targets currently marketed or in clinical trials (NGF, PCSK9, CD19, CD20, PD1 and CTLA4) and a set of in-house targets of interest (LDHA, cAbl, DHFR). Target engagement was observed for the cAbl-ThermLuc fusion using both dasatinib and GNF-2, which are orthosteric and allosteric inhibitors, respectively (Supplementary Fig. S4B). We also explored DHFR thermal melt and target engagement using the RT-CETSA system, which revealed a potential caveat of attaching a thermally stable luciferase to a target-of-interest. For DHFR, we observed a thermal stabilization of the ThermLuc fusion, where a partial aggregation profile was observed (Supplementary Fig. S4C). The thermal shift induced by the ligand methotrexate was detectable for DHFR-ThermLuc, but smaller in magnitude compared to DHFR-SplitLuc.

We further hypothesized the ThermLuc effects on DHFR thermal stability could be altered by varying the peptide linker between the two proteins. A set of 17 linkers with a range of predicted rigidity were constructed. Rigid polyproline-containing linkers further increased thermal stability of the DHFR-ThermLuc fusion protein. Some linkers reduced the overall thermal stability of the fusion protein however none fully recapitulated the melting behavior or thermal shift observed for DHFR with the small SplitLuc peptide tag (Supplementary Fig. S4D, E). Additional research is needed to define intermolecular effects of ThermLuc on targets and its propensity to alter the inherent stability of its fusion partner. Our proof-of-concept studies indicates that RT-CETSA has the potential to serve as a multi-target platform that can rapidly assess EC_50_ values across a variety of different targets in parallel and can be used for SAR studies in medicinal chemistry campaigns.

## Discussion

The practicality of RT-CETSA as a high-throughput target engagement method relies on its ability to monitor target unfolding across a temperature range inside a living cell. Here we showed the first reported examples of recording the entire melting profile of a target, within cells, in real time. We also showcased the first instrument capable of recording luminescence during heating and the first examples of using thermally stable luciferase fusion proteins to assess target engagement for a variety of targets on the same plate. The RT-CETSA platform was shown to successfully characterize target engagement and support SAR assessment using a set of 29 LDHA_i_ analogs with EC_50_ values comparable to other binding assays.

Demonstrating on-target activity in cell-based models is a critical step in the development of small molecule probes and therapeutic candidates. Downstream validation of target engagement can be especially challenging when switching from biochemical assays to physiologically relevant cellular models. Many preclinical candidates fail because of off-target effects or poor physicochemical and pharmacokinetic properties (28–30). This is exemplified within a set of LDHA inhibitors, where potency and number of active analogs decreased as the complexity of the target’s microenvironment increased from purified protein to cellular models (biochemical enzymatic assay vs. cell-based lactate assay; SPR and DSF vs. CETSA). For example, compound **25** had a biochemical enzymatic assay IC_50_ value of 300 nM but its potency diminished in a cell-based lactate assay with an IC_50_ value of 5 μM. A common explanation is that biochemical, SPR and DSF binding assays use recombinant protein and may overestimate the capacity of a molecule to engage a target within cells. Furthermore, cell-based phenotypic assays, such as measuring lactate levels, can be susceptible to false positives driven by off-target activity which result in overlapping phenotypes.

CETSA has become a valuable and widely implemented method for assessing target engagement in a cellular environment. All known CETSA methods involve endpoint lytic detection and either require high affinity antibodies, or have a reliance on luciferase reporters like NLuc that can alter target biology and drive aggregation at temperatures lower than a target’s inherent thermal properties (13, 23, 24, 31, 32). The thermal and mechanical instability of the small and highly luminescent NLuc protein could therefore impede the high-throughput transformation of CETSA for some targets with high a T_agg_ (24, 33). These issues led to the conceptual design and bioengineering of more thermally stable ThermLuc fusions as a reporter for RT-CETSA.

By monitoring the entire thermal profile for each sample, RT-CETSA enables multiplexing of multiple targets, compounds, and concentrations within a single experiment. This circumvents optimizing for specific melting temperatures and removes the risk of selecting a temperature where the thermal shift window would be missed or masked in a traditional CETSA experiment. Consistently, RT-CETSA analysis of LDHA inhibitors showed concentration-dependent ligand-induced thermal shifts with EC_50_ values that were highly correlated with previous CETSA data (endogenous and SplitLuc CETSA) as well as biophysical target engagement data. However, the absolute potencies were not identical across the assays. This is not surprising. Several studies have demonstrated that potency in isothermal CETSA is highly dependent on experimental conditions including duration of the heating step (34, 35).

In our RT-CETSA studies, we found that longer heating times or higher temperatures decreased the apparent potency for stabilization. Our SplitLuc CETSA data on the 29 LDHA_i_ analogs is consistent with this interpretation, where heating at 69 °C uniformly right shifted and showed diminished potency compared to heating at 65 °C or 61 °C. These results further highlight the significant risk of missing target engagement events when using single endpoint recordings if a non-optimal temperature is selected. It also highlights a need to improve analysis methods and utilize metrics beyond the apparent T_agg_. Therefore, we developed a novel method of analysis to pair with the RT-CETSA technology by modifying previously reported thermal proteome profiling analysis methods known as NPARC (nonparametric analysis of response curves). The modified NPARC was more sensitive and specific to ligand-induced thermal stabilization than calculating a melting point or other single parameter values (22, 36).

Correlative studies comparing RT-CETSA analysis methods with other binding assays, like DSF and SPR, revealed that NPARC showed >95% overlap in identified “hits” compared to classical methods of CETSA analysis using AUC and T_agg_. By comparing goodness-of-fit tests for dose-response data across the entire melting curve, we captured treatment stabilization effects that were not detected by single summary statistics commonly used to describe thermal unfolding data. Minimum significance ratio (MSR) and Z’ assay reproducibility calculations, two widely used measures of assay quality that describes separation between positive and negative controls (37), both supported the modified NPARC analysis as a more robust and reproducible analysis method compared to T_agg_ and AUC. Moreover, SAR analysis of a set of LDHA inhibitors using the NPARC method supported previous studies that defined the importance of an ethyne linker between phenyl pyrazole and thiophene, or similar bioisosteric rings, as important for elevated intracellular and *in vivo* inhibitory activity (27, 38). This highlights the reliability of RT-CETSA to quantitatively determine SAR between sets of active and inactive analogues.

RT-CETSA bridges the gap between high data content methods that are traditionally low-throughput with more accessible high-throughput but single parameter target engagement methods, and couples these technologies in living cells. Developing the RT-CETSA technology foremost required the creation of thermally stable ThermLuc reporters and building a prototype instrument with a core interface that couples a high-precision, high-speed PCR block with a sensitive CCD system capable of capturing luminescent signals continuously throughout the thermal ramp. There is currently no instrument on the market today that pairs these two components together. To validate the RT-CETSA method, we chose an existing PCR platform, the Roche LightCycler 480 II, to serve as the heating platform. Preliminary testing showed that the OEM instrument was insensitive to luminescence signals that are in the working range of commonly used microplate readers (ViewLux, Tecan). Therefore, we built a prototype machine by removing the xenon bulb and all emission filters from the instrument light path and swapped the existing CCD detection unit out for a more sensitive camera capable of recording luminescence, the Hamamatsu Orca R2.

The entire melting profiles of a variety of targets were capably visualized and recorded from the same plate, despite the small signal window of luminescence below saturation but above noise from our crudely assembled prototype. In a typical RT-CETSA experiment, the entire thermal melt profile of a target inside a live cell can be recorded in less than 4 min. 55 readings at discrete temperatures are recorded from each well in 4 sec intervals with Δ1 °C temperature increments. RT-CETSA showed that thermal unfolding rapidly occurs upon the application of heat and stable equilibrium of fraction unfolding is reached within 4-8 sec of an isothermal temperature hold. Previous work on standardizing differential scanning fluorimetry protocols noted a phenomenon where T_m_ readouts are highly dependent on experimental parameters like ramp speed and temperature holds, but the ΔT_m_ induced by ligand binding remained constant (39). Increasing the hold time of each temperature to 20 sec reduced the T_agg_ of LDHA by Δ−16 °C, but the increased hold time did not significantly impact the compound-induced thermal shift. 4 sec holds at each temperature was sufficient to capture enough signal using the prototype detection system while also limiting the decay rate of the furimazine substrate that would occur with longer holds. We expect that these hold times could be reduced further with a more sensitive camera, which would allow for faster ramping through a range of temperatures.

The quick workflow and limited hands-on requirements of the RT-CETSA method lend to a quick adaptation into a variety of applications, including characterizing chemical probes, lead optimization, identification of allosteric binders or PROTACs, and/or probing against multiple targets (e.g. family of proteins, alternate pathways). The prototype instrument enabled us to assess the melt profile of nine different protein targets simultaneously despite differences in starting luminescence, functional activity, and subcellular localization. We also demonstrated the compatibility of RT-CETSA to detect the melting profiles of secreted PCSK9 and NGF1 protein targets with no prior purification or enrichment of target needed. We anticipate entire families of protein could be rapidly assessed for ligand binding under identical cellular conditions and provide valuable insights into off-target binding for sets of compounds. RT-CETSA may be particularly valuable for cellular proteins that are hard to ‘extract’, such as nuclear proteins, using CETSA-compatible methods under non-denaturing conditions (40). The RT-CETSA method has potential to capture the melting behavior of proteins in various subcellular compartments, provided that furimazine can access the target-ThermLuc fusion.

Importantly, RT-CETSA is prone to its own limitations and caveats, some of which are shared with traditional CETSA. For instance, some compounds that engage with a target do not affect its thermal stability. Additionally, cellular membrane permeability is disrupted at temperatures greater than 60 °C, which can confound interpretations at high melting temperatures (17, 41, 42). Similarly, higher temperatures are expected to alter the permeability of the furimazine substrate and enzymatic activity of the ThermLuc luciferase. Finally, our studies with DHFR indicate that ThermLuc may alter the melting behavior of some targets. We believe the impact of these factors can be minimized with proper controls and orthogonal screening methods. In summary, RT-CETSA provides an adaptable method that is broadly applicable for target engagement and screening campaigns, while offering sensitivity and ease-of-use that are unparalleled by current CETSA methods.

## Acknowledgements

We thank the NCATS compound management group for sourcing, quality control, formatting, and plating of compounds. We also thank Kyle Brimacombe for contributions to graphics.

## Author Contributions

T.W.S., M.H.R., A.E.O., N.S., and M.J.H. contributed to the bioengineering, experimentation, development, and analysis of the RT-CETSA datasets and methodology. T.W.S., M.H.R., A.E.O., K.B., G.R., J.J.M., S.M., B.B., A.S. and M.J.H. all contributed to the conceptual design of the RT-CETSA technology and platform. T.W.S., M.H.R., A.E.O., A.M., D.T., B.B., N.S., and M.J.H. contributed to the writing, figure generation, and editing of the manuscript. T.V. contributed to the creation of the MATLAB script used to analyze the luminescence signal. T.W.S., M.H.R., E.W., S.M., M.J.H. contributed to the creation and optimization of the prototype RT-CETSA machine and analysis software. D.T. and G.R. contributed to the synthesis of the furimazine substrate and LDHA inhibitors.

## Disclosures

T.W.S., M.H.R., A.E.O., B.B., S.M., A.S. and M.J.H. are inventors on a patent REAL-TIME CELLULAR THERMAL SHIFT ASSAY (RT-CETSA) FOR RESEARCH AND DRUG DISCOVERY (PCT/US21/45184). T.W.S., M.H.R., B.B. N.S., S.M. A.S., T.V., and M.J.H. are inventors on a related provisional patent METHODS AND SYSTEMS FOR ANALYZING TARGET ENGAGEMENT DATA FROM BIOLOGICAL ASSAYS (HHS E-022-2022-0-US-01).

## Methods

### Real-time CETSA assay

The RT-CETSA procedure involves transfecting cells with a plasmid encoding the target fused to ThermLuc and heating the sample while monitoring luminescence in real-time. Briefly, 5000 cells expressing the TOI-ThermLuc fusion protein were dispensed into 384-well PCR plates (10 μL per well) in CETSA buffer (phenol-free high glucose DMEM with sodium pyruvate and 1X Glutamax without FBS). 20 nL of compounds or DMSO vehicle controls were acoustically dispensed into the cells and the plates were incubated for 1 hour at 37 °C. 10 μL of 2X furimazine (diluted from Promega 50X stock into CETSA buffer) solution was added to the 10 μL of transfected HEK293T cells in each well. The plate was sealed then run on a modified Roche LightCycler 480 II with luminescence recorded kinetically as temperature increased stepwise (e.g. 1 °C increments from 37 °C to 90 °C) to define the melt profile. An optimal density was determined to ensure that the signal during heating would remain in the camera’s linear detection range. The images taken were processed with an in-house MATLAB script to extract the raw luminescence values from each well at each temperature.

### Prototype RT-CETSA hardware

All emission filters were removed from the light path in the Roche LightCycler 480 II for RT-CETSA experiments. After instrument warmup and pre-run checks with the stock camera in place to allow the machine to initialize, the xenon bulb in the light cycler was removed and covered to shield the plate from excess light and the original LightCycler camera was swapped with a Hamamatsu Orca II camera. It was important that the instrument be left in its stock configuration upon power-up so that pre-run checks could proceed without error. The Roche LightCycler control software was set to run a protocol including a 2 min hold at 37 °C to allow for camera equipment swaps and enclosure reset after the instrument had completed pre-run checks with the stock camera in place. Once the equipment changes were made and the 2 min pre-run hold ended, LabView Software interfaced with the Orca II camera took exposures at user-defined intervals until the operator stopped operation at the end of the heating cycle. The temperature ramp was set so that the time to ramp and the time held at each step/°C in the thermal ramp was equal to the shutter speed of the camera (in testing, was either 2, 4, or 8 sec).

### Melting curve analysis

Raw images from the RT-CETSA platform were run through a MATLAB script to obtain luminescence values for each well across each image. Digital luminescence images were analyzed using a customized MATLAB (The Mathworks, Inc.) script (public file source). The automated workflow is briefly described in the following section. For time-series analysis of RT-CETSA images, the calibration steps (details below) are required only on the initial time point. All subsequent time-series images were automatically processed in batch mode, and results for each sample were exported as time-indexed series tables. Prior to running the analysis, the user defines the script parameters to account for the size of the square bounding box region surrounding each sample in the image (estimated number of pixels, width, and height). When executed in the MATLAB computational environment (MATLAB version 2021a) the script functions allow the user to select the first image file in the time series for display, and a graphical user interface prompts the user to locate the positions of the upper left sample, the upper right sample, and the lower left sample. The user also enters the total number of sample rows and columns that are in the selected contiguous block of samples. Based on the above calibration information, the script functions adjust the image to correct for any rotation of the samples and applies an evenly spaced analysis grid. Each grid region contains luminescence information from a single sample. An automatic threshold is applied to each region to segment the signal region from the local background region within each grid region. The size (in pixels) of the signal region is reported and can be automatically gated to eliminate irregular signals (for example, small false positive speckle noise regions or large false positive regions when no true signal region is present in the grid). Numerical values for mean signal intensity per signal region, mean local background intensity, and mean pixel intensity region minus local background are reported for each analyzed grid region. These raw values were then processed using MoltenProt (CSSB, Hamburg Germany) software using the standard two-state unfolding model with 15 °C of the beginning and ending of curves used as initial values for baseline fit estimates. T_agg_ values were derived as the midpoint of the 4-parameter sigmoidal curve fit of a baseline corrected thermal unfolding curve. AUC values were calculated by processing the baseline-corrected curve fits using the auc function in the R package ‘MESS’. All dose-response fits were calculated using a 4-parameter log-logistic fit using the R package ‘drc’ with 10000 iterations to convergence. Nonparametric analysis of the thermal curves was performed by fitting two models to each dose-response at each data collection temperature point: 1) null model, which is a linear fit with a slope of 0, and 2) alternate model, which is a 4-parameter log-logistic fit. The residual sum of squares was then calculated for each model across every temperature point, and a nonparametric Mann Whitney U test against model RSS values were performed to determine significant stabilization. EC_50_ values were then derived from curves that have significant stabilizers by fitting a 4-parameter log-logistic fit using the curve values at the point of maximal RSS difference between null and alternate models (9, 20, 26, 33).

### Molecular biology

ThermLuc plasmids were created using a pcDNA3.1(+) backbone and cloning into the NheI and EcoRI sites using Genestrands (Eurofins) encoding LgBiT-[1/3/6/9/12/15 GlySer]-HiBiT-GlySer. The gene strand was inserted into the linearized backbone using InFusion (Takara) following the manufacturer’s instructions. For TOI-ThermLuc fusions, the sequence ggatccGGCGGTGGTGGCTCT (GlySerGlyGlyGlyGlySer) was placed immediately upstream of ThermLuc, to create a BamHI (in lowercase above) site such that various targets could be cloned as N-terminal fusions using NheI and BamHI sites and InFusion reagents. For transmembrane and secreted targets (CD19, CD20, NGF, PCSK9), fusions were created in TOI-ThermLuc orientation to maintain signal sequences and partitioning to the secretory pathway. An acceptor plasmid containing ThermLuc in the N-terminal orientation was created by cloning the ThermLuc gene strand into pcDNA3.1(+) using the NheI and EcoRI sites, where the sequence GGCGGTGGTggatcc (GlyGlyGlyGlySer) was placed immediately downstream of ThermLuc to create a BamHI site (in lowercase above) such that various targets could be cloned as C-terminal fusions using BamHI and EcoRI sites and InFusion reagents. To create plasmids encoding DHFR-ThermLuc with different intervening linkers, we first created a pcDNA3.1(+) construct in which the nucleotide sequence ggatccGCTGAAGCCGCGGCTAAAGAGGCTGCCGCGAAAGAAGCTGCAGCTAAGGAGGCTGCAGCGAAAGAGGCAGCGGCAAAGGAGGCTGCCGCGAAGGCTaagctt inserted between the final amino acid of DHFR and the start codon of ThermLuc. This represented DHFR-A[EAAAK]_6_A and allowed subsequent digestion with BamHI and HindIII (sites in lowercase in sequence above) and insertion of linker sequences using InFusion PCR. Linker sequences were synthesized as two complementary oligonucleotides (Supplementary Table 1) that were annealed in a 50 μL solution containing 10 μM forward oligo, 10 μM reverse oligo, 10 mM TrisHCl (pH 8.0), 50 mM NaCl and 1 mM EDTA. The solution was placed in a beaker of boiling water, beaker was removed from heat, and tube remained in the water to cool to room temperature without intervention. The resulting duplex was used for InFusion PCR following the manufacturer’s instructions.

### Cell culture

HEK293T cells were grown in a high-glucose DMEM with sodium pyruvate (Gibco) plus 10% FBS (Hyclone), 1X GlutaMax (Gibco), 100U/ml penicillin, and 100U/ml streptomycin. Cells were incubated at 37 °C with 5% CO_2_ and 95% humidity. Cells were routinely tested for mycoplasma using a Lonza MycoAlert kit.

### Chemistry

All compounds were dissolved in DMSO. LDHA inhibitors were titrated in two-fold serial dilutions and 20 nL were transferred using acoustic dispenser Echo 550 (Labcyte). DMSO concentrations were maintained at less than 0.5% total volume. Furimazine was synthesized in-house as previously reported and stored at −20 °C as a solid powder (43).

### Endpoint CETSA protocol to test ThermLuc fusions

HEK293T cells were transiently transfected with plasmids containing TOI-luciferase variants using a reverse transfection procedure. Briefly, 3 μg of the plasmid and 6.25 μl of Lipofectamine 2000 was combined in 1.25 mL Opti-MEM (Gibco) to create the transfection complex. Following a 20-min incubation, the transfection complex was added to 1.25 x 10^6^ cells suspended in 1.25 mL of a phenol-free high glucose DMEM with 10% FBS and 1X GlutaMax in a six-well plate. After 24 or 48 hours, cells were lifted with 0.25% trypsin and resuspended in a phenol-free high glucose DMEM with 1X GlutaMax without FBS. Next, 10 μl of cells per well were dispensed into white 384-well PCR plates (Roche) using a Multidrop Combi (ThermoFisher Scientific). Next, an Echo 550 was used to acoustically transfer DMSO or compound to the cells, which were then incubated for an hour at 37 °C with CO_2_ and 95% humidity. Following the incubation, cells were heated for 3.5 min using a 384-well thermal cycler; then 10 μL of 2X furimazine (Promega 50X stock) solution diluted in phenol-free high glucose DMEM with GlutaMax was added to each well.

For a 96-well plate, cells were treated in bulk with DMSO or compound then transferred to PCR tubes (30 μL/ tube) for subsequent incubation and heating. Once cooled to room temperature, 10 μL of cells from each tube was transferred to a white 384 well plate then 10 μL of 2X furimazine was added to each well. Luminescence was detected using a PerkinElmer ViewLux microplate reader equipped with clear filters.

### SplitLuc CETSA

Cells transfected with LDHA fused to an 86b peptide were aliquoted for CETSA experiments as previously described (17). Briefly, after 24h transfection cells were harvested by trypsinization, resuspended at 1 million cells per mL in CETSA buffer and dispensed (10 μL per well) into 384-well PCR plates (Roche) using a Multidrop Combi (ThermoFisher). 20 nL of compounds or DMSO vehicle controls were acoustically transferred using an Echo 550 and incubated for 1 hour at 37 °C. Plates were sealed and heated at 37, 61, 65 and 69 °C for 3.5min and cooled to 25 °C using qPCR machine (Applied Biosystems) using ramp speed of 1.5 °C/sec for heating phase and max ramp rate for the cooling phase. 2 μL of 6% NP40 were added per well and incubated at room temperature for 30 min to allow cell lysis, followed by the addition of 11S and furimazine substrate at final concentrations of 100nM and 0.5X, respectively. Samples were analyzed for luminescence intensity using a ViewLux reader equipped with clear filters (Perkin Elmer).

### NanoDSF

The interaction between LDHA and a small molecule was evaluated using a label-free approach. Specifically, 30 μL of 0.1 mg/mL recombinant LDHA in assay buffer [25 mM Tris, 100 mM NaCl, pH 7.5] was incubated with 50 μM compound for 10 min at room temperature, and then loaded into 3 standard capillaries for triplicate readings in a Prometheus NT.48 instrument (Nanotemper Technologies, Munich, Germany). Experiments were run using 40% excitation power, and a 1° C/min temperature ramp. Analysis was performed on the instrument using manufacturer’s software to derive the temperature of melting and first derivatives for each sample. The experiments determining the thermal stabilization between LgBiT and HiBiT were performed as described above. LargeBiT (11S) large fragment and HiBiT peptide were synthesized as previously described and diluted to 14 μM and 30 μM in PBS, respectively (17). The 156 and NP (native peptide) proteins consist of the first 156 amino acids and final 13 amino acids, respectively, of unmodified NanoLuc (23) (Genscript).

### Cellular lactate production assay

HEK293T cells were cultured as described above. Cells were trypsinized and resuspended in phenol red free DMEM (Life Technologies) without supplements. Cells were immediately plated to 1536-well black clear bottom plates (Corning) at 250 cells per well in 4 μL volume. Compound or vehicle control was added to wells via pin tool transfer and cells were incubated at 37 °C for 1 h. Two μL of lactate reaction mixture (Biovision K607-100) was added to each well and plates were covered and incubated at room temperature for 30 min. Fluorescence was measured using a ViewLux microplate imager equipped with Ex/Em 528/598nm filters.

### Cellular viability assay using Cell Titer glo

CellTiter-Glo (Promega) experiments were conducted according to the manufacturer’s protocol. Briefly, 5000 cells were dispensed into 384-well plates (10 μL per well) and treated with 20 nL of LDHA inhibitors and DMSO controls for 1 hour at 37 °C. 10 μL of CellTiter-Glo reagent was then added and the plate was incubated at room temperature for 30 min with continuous shaking. Luminescence was detected using a PerkinElmer ViewLux microplate reader equipped with clear filters.

### SYSTEMETRIC Cell Health Screen

The Cell Health Screen (AsedaSciences) is a multiparametric acute cell stress assay using a panel of fluorescent physiological reporting dyes on an automated flow cytometry platform with a supervised machine learning (ML) classifier using a multiparametric logistic model (44). The final probability score, or “Cell Health Index”, is a quantitative assessment of a multiparametric phenotype’s similarity to a diverse set of known bad actors. In a 384-well platform, HL60 cells (100,000 cells in 40 μL volume) were exposed to a 10-step, 3X dilution series of each test compound (5nM – 100 μM) for 4 hours. After compound exposure, cells were stained with a panel of fluorescent dyes that report physiological signatures of both mitochondrial dysfunction and gross cell stress. Fluorescence data were collected using automated flow cytometry, still in the original 384-well plate, with no gating. In addition, forward scatter and side scatter at 488nm were acquired for conversion into a cell morphology parameter. For each test compound, ungated detection parameters were converted to a tensor of values based upon QF distances between each step in the dilution series and both the positive and negative control wells in each row of the assay plate. This tensor becomes the input used for supervised machine learning classification relative to the training set of 300 known compound drawn from on-market pharmaceuticals, withdrawn drugs, research compounds, and a few industrial/agricultural compounds. First, all training set compounds were assigned to either the “positive” or “negative” training class based upon external information from the scientific literature, clinical trial reports, and/or known commercial histories. The classifier was trained by first dividing the training set into the two training classes, based upon external information, and then optimizing the fit of its multiparametric logistic model on the empirical screening data for all training compounds. The training set contained an approximate 1:3 ratio of “high cell stress” to “low cell stress” screen phenotypes, defined by an arbitrary cutoff at a probability of 0.5 that the classifier can assign any individual compound to the “high cell stress” class based upon the similarity of its screen phenotype to the rest of that class. For each test compound, the final multiparametric risk score, or Cell Health Index (CHI), is the probability with which the test compound’s screen phenotype can be assigned to the “high cell stress” class defined by the training set. In addition, a unidimensional version of the classifier was applied to the detection parameters separately, calculating probability of “high cell stress” class assignment if only data for that parameter are considered.

## References

1. Vidal M, Cusick Michael E, & Barabási A-L (2011) Interactome Networks and Human Disease. Cell 144(6):986–998.

2. Ciulli A (2013) Biophysical screening for the discovery of small-molecule ligands. Methods Mol Biol 1008:357–388.

3. Edink E, et al. (2011) Fragment growing induces conformational changes in acetylcholine-binding protein: a structural and thermodynamic analysis. J Am Chem Soc 133(14):5363–5371.

4. Roy MJ, et al. (2019) SPR-Measured Dissociation Kinetics of PROTAC Ternary Complexes Influence Target Degradation Rate. ACS Chem Biol 14(3):361–368.

5. Fedorov O, Niesen FH, & Knapp S (2012) Kinase inhibitor selectivity profiling using differential scanning fluorimetry. Methods Mol Biol 795:109–118.

6. Layton CJ & Hellinga HW (2010) Thermodynamic Analysis of Ligand-Induced Changes in Protein Thermal Unfolding Applied to High-Throughput Determination of Ligand Affinities with Extrinsic Fluorescent Dyes. Biochemistry 49(51):10831–10841.

7. Niesen FH, Berglund H, & Vedadi M (2007) The use of differential scanning fluorimetry to detect ligand interactions that promote protein stability. Nature Protocols 2:2212–2221.

8. Jafari R, et al. (2014) The cellular thermal shift assay for evaluating drug target interactions in cells. Nat Protoc 9(9):2100–2122.

9. Molina DM, et al. (2013) Monitoring Drug Target Engagement in Cells and Tissues Using the Cellular Thermal Shift Assay. Science 341(6141):84–87.

10. Mateus A, et al. (2020) Thermal proteome profiling for interrogating protein interactions. Mol Syst Biol 16(3):e9232.

11. Franken H, et al. (2015) Thermal proteome profiling for unbiased identification of direct and indirect drug targets using multiplexed quantitative mass spectrometry. Nat Protoc 10(10):1567–1593.

12. Friman T (2020) Mass spectrometry-based Cellular Thermal Shift Assay (CETSA(R)) for target deconvolution in phenotypic drug discovery. Bioorg Med Chem 28(1):115174.

13. Mateus A, et al. (2020) The functional proteome landscape of Escherichia coli. Nature 588(7838):473–478.

14. Henderson MJ, Holbert MA, Simeonov A, & Kallal LA (2020) High-Throughput Cellular Thermal Shift Assays in Research and Drug Discovery. SLAS Discov 25(2):137–147.

15. Kubota K, Funabashi M, & Ogura Y (2019) Target deconvolution from phenotype-based drug discovery by using chemical proteomics approaches. Biochim Biophys Acta Proteins Proteom 1867(1):22–27.

16. Robers MB, et al. (2015) Target engagement and drug residence time can be observed in living cells with BRET. Nat Commun 6:10091.

17. Martinez NJ, et al. (2018) A widely-applicable high-throughput cellular thermal shift assay (CETSA) using split Nano Luciferase. Scientific reports 8(1):9472–9472.

18. Dart ML, et al. (2018) Homogeneous Assay for Target Engagement Utilizing Bioluminescent Thermal Shift. ACS Med Chem Lett 9(6):546–551.

19. Seashore-Ludlow B, Axelsson H, & Lundback T (2020) Perspective on CETSA Literature: Toward More Quantitative Data Interpretation. SLAS Discov 25(2):118–126.

20. Massey AJ (2018) A high content, high throughput cellular thermal stability assay for measuring drug-target engagement in living cells. PLoS One 13(4):e0195050.

21. Shaw J, et al. (2018) Determining direct binders of the Androgen Receptor using a high-throughput Cellular Thermal Shift Assay. Sci Rep 8(1):163.

22. Childs D, et al. (2019) Nonparametric Analysis of Thermal Proteome Profiles Reveals Novel Drug-binding Proteins. Mol Cell Proteomics 18(12):2506–2515.

23. Dixon AS, et al. (2016) NanoLuc Complementation Reporter Optimized for Accurate Measurement of Protein Interactions in Cells. ACS Chem Biol 11(2):400–408.

24. Hall MP, et al. (2012) Engineered luciferase reporter from a deep sea shrimp utilizing a novel imidazopyrazinone substrate. ACS Chem Biol 7(11):1848–1857.

25. Sanchez TW, et al. (2021) High-Throughput Detection of Ligand-Protein Binding Using a SplitLuc Cellular Thermal Shift Assay. Methods Mol Biol 2365:21–41.

26. Kotov V, et al. (2021) In-depth interrogation of protein thermal unfolding data with MoltenProt. Protein Science 30(1):201–217.

27. Rai G, et al. (2017) Discovery and Optimization of Potent, Cell-Active Pyrazole-Based Inhibitors of Lactate Dehydrogenase (LDH). J Med Chem 60(22):9184–9204.

28. Morgan P, et al. (2012) Can the flow of medicines be improved? Fundamental pharmacokinetic and pharmacological principles toward improving Phase II survival. Drug Discov Today 17(9-10):419–424.

29. Vinegoni C, et al. (2015) Advances in measuring single-cell pharmacology in vivo. Drug Discov Today 20(9):1087–1092.

30. Walker DK (2004) The use of pharmacokinetic and pharmacodynamic data in the assessment of drug safety in early drug development. Br J Clin Pharmacol 58(6):601–608.

31. Nagasawa I, et al. (2020) Identification of a Small Compound Targeting PKM2-Regulated Signaling Using 2D Gel Electrophoresis-Based Proteome-wide CETSA. Cell Chem Biol 27(2):186–196 e184.

32. Savitski MM, et al. (2014) Tracking cancer drugs in living cells by thermal profiling of the proteome. Science 346(6205):1255784.

33. Ding Y, Apostolidou D, & Marszalek P (2020) Mechanical Stability of a Small, Highly-Luminescent Engineered Protein NanoLuc. Int J Mol Sci 22(1).

34. Jarzab A, et al. (2020) Meltome atlas-thermal proteome stability across the tree of life. Nat Methods 17(5):495–503.

35. Maatta TA, et al. (2020) Aggregation and disaggregation features of the human proteome. Mol Syst Biol 16(10):e9500.

36. Perrin J, et al. (2020) Identifying drug targets in tissues and whole blood with thermal-shift profiling. Nat Biotechnol 38(3):303–308.

37. Zhang JH, Chung TD, & Oldenburg KR (1999) A Simple Statistical Parameter for Use in Evaluation and Validation of High Throughput Screening Assays. J Biomol Screen 4(2):67–73.

38. Rai G, et al. (2020) Pyrazole-Based Lactate Dehydrogenase Inhibitors with Optimized Cell Activity and Pharmacokinetic Properties. J Med Chem 63(19):10984–11011.

39. Baljinnyam B, Ronzetti M, Yasgar A, & Simeonov A (2020) Applications of Differential Scanning Fluorometry and Related Technologies in Characterization of Protein-Ligand Interactions. Methods Mol Biol 2089:47–68.

40. Scheer U, Kartenbeck J, Trendelenburg MF, Stadler J, & Franke WW (1976) Experimental disintegration of the nuclear envelope. Evidence for pore-connecting fibrils. J Cell Biol 69(1):1–18.

41. Bischof JC, et al. (1995) Dynamics of cell membrane permeability changes at supraphysiological temperatures. Biophys J 68(6):2608–2614.

42. McNulty DE, et al. (2018) A High-Throughput Dose-Response Cellular Thermal Shift Assay for Rapid Screening of Drug Target Engagement in Living Cells, Exemplified Using SMYD3 and IDO1. SLAS Discov 23(1):34–46.

43. Shakhmin A, et al. (2016) Three Efficient Methods for Preparation of Coelenterazine Analogues. Chemistry 22(30): 10369–10375.

44. Bieberich AA, et al. (2021) Acute cell stress screen with supervised machine learning predicts cytotoxicity of excipients. J Pharmacol Toxicol Methods 111:107088.

