## Supplementary Figures S1-S4 for "A real-time cellular thermal shift assay (RT-CETSA) to monitor target engagement"

**Supplementary Material Contents:**

Figures S1-S4

Supplementary Movie Legend

Supplementary Figure S1

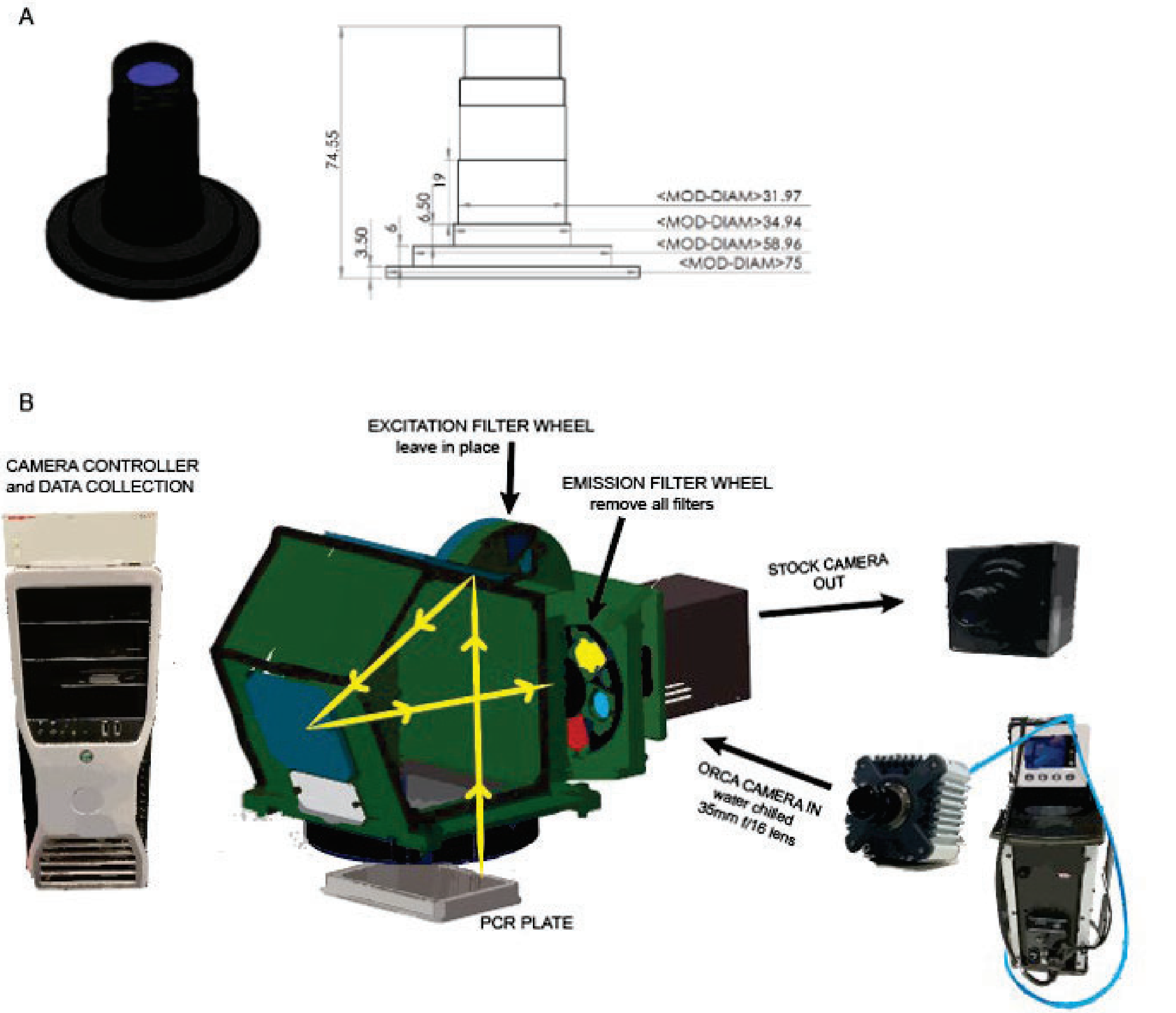


*Fig. S1: RT-CETSA prototype device. **(A)** OEM camera on the Roche LC480 II Real-Time PCR instrument. Dimensions of the stock CCD lens, which are held within a metal bracket in the machine, are shown. **(B)** Schematic of RT-CETSA components. Emission filters were removed and the stock camera was replaced with a water-chilled Orca R2 CCD camera for an unfiltered light path and improved luminescence detection. The 35mm f/1.6 lens was a direct replacement for the original stock CCD lens aperture and focal length.*

Supplementary Figure S2

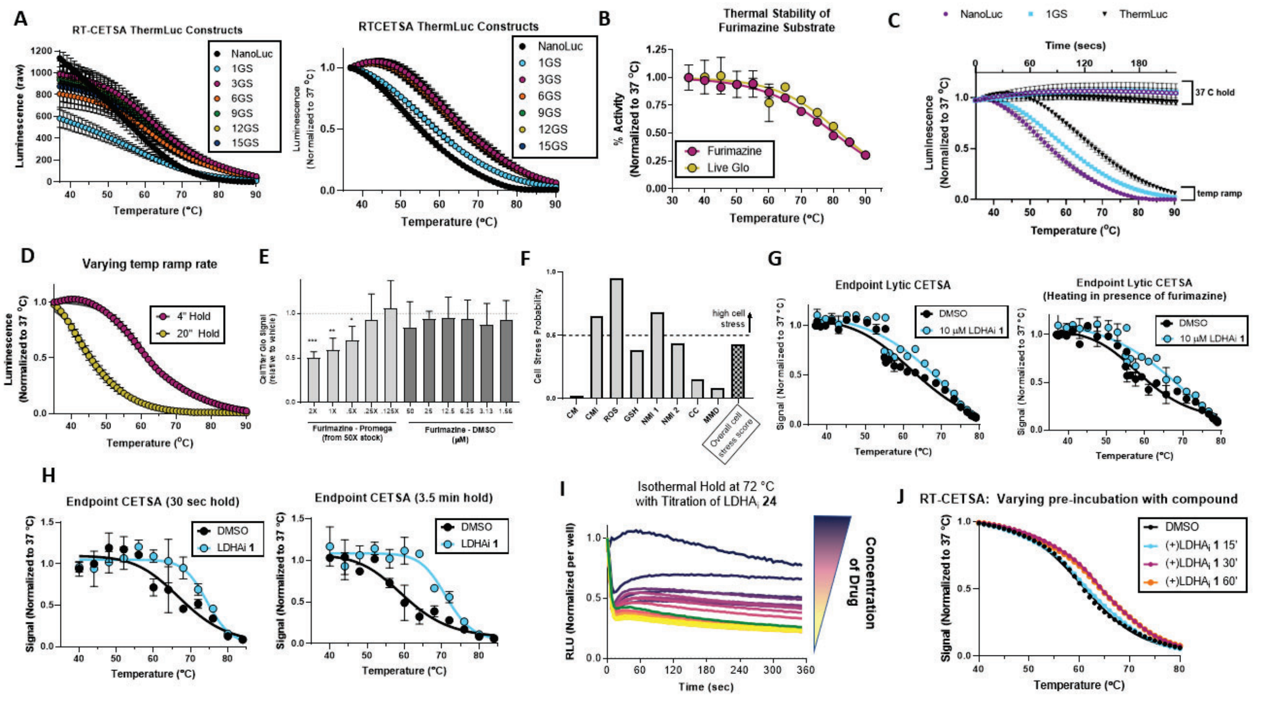


*Fig S2: **(A)** Thermal melt profile of NLuc variants with different Gly-Ser linker lengths between the 11S and HiBiT fragments, as measured using RT-CETSA device (mean ± SD, n=64 wells) and a commercially available furimazine (NanoGlo, Promega). Right panel depicts data normalized to starting luminescence for each individual well. **(B)** Thermal stability of furimazine (NanoGlo, Promega) and LiveGlo (Promega) substrates were assessed by first heating 0.2X substrate to the indicated temperature for 3.5min in the presence of HEK293 cells (untransfected), cooling to room temperature, and then adding recombinant 11S and HiBiT peptide. Luminescence was measured (mean ± SD, n=4). **(C)** Thermal stability of furimazine synthesized in-house was examined in the RT-CETSA device, either with temperature ramp or using a 37 $^o$C hold, for identical timeframes (n=64 wells). **(D)** Melting profile of LDHA-ThermLuc was examined using different temperature ramping, either holding at each temperature integer for 4 sec or 20 sec (mean ± SD, n=96 wells). **(E)** Cell viability after 48h incubation with furimazine, diluted from Promega 50X stock or DMSO solution (mean ± SD, n=4). *p<0.05, **p<0.01, ***p<0.001. **(F)** Systemetric CellHealth screening performed using furimazine formulated in DMSO. CM= cell morphology; CMI=cell membrane integrity; ROS = reactive oxygen species; GSH = glutathione; NMI1/2 = nuclear membrane integrity metric 1 and 2; CC = cell cycle; MMD = mitochondrial membrane depolarization. Overall score of 0.4 is threshold for high cell stress. **(G)** Target engagement and thermal shift of LDHA-ThermLuc was examined using endpoint CETSA analysis, where furimazine was excluded (left) or included (right) during the heating step. Fresh substrate was added to all samples before measuring luminescence (mean ± SD, n=2). **(H)** Thermal shift was examined for LHDA-ThermLuc using the endpoint method after treatment with 10 µM LDHAi **63**. Samples were heated for 30 sec (left) or 3.5 min (right) before lysis. Furimazine was added and luminescence was measured (mean ± SD, n=2). **(I)** Rapid aggregation of LDHA-ThermLuc after transitioning to 72 °C. Luminescence was monitored using the RT-CETSA device. **(J)** RT-CETSA analysis of LDHA-ThermLuc thermal shift for cells pre-treated with 10 µM LDHA$_i$ **63** for 15, 30, or 60 min.*

Supplementary Figure S3

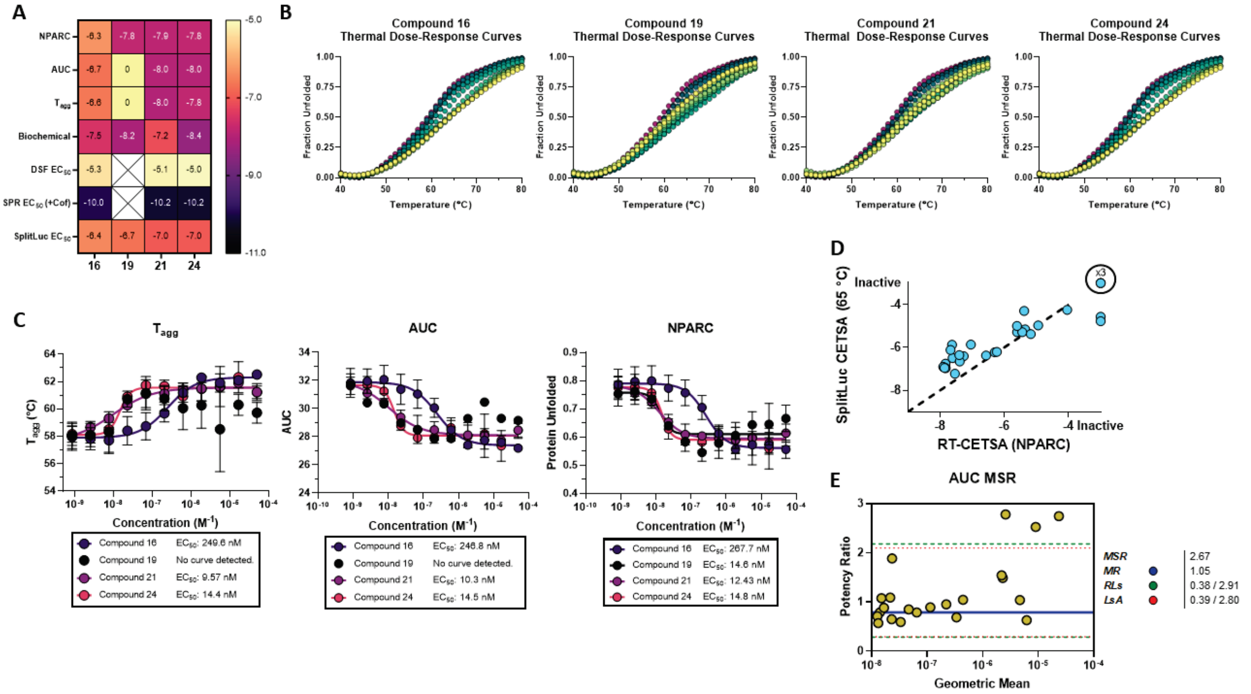


*Fig S3* ***(A)*** *Log $EC_{50}$ values for highlighted prior art compounds when analyzed with RT-CETSA methods, acoustic CETSA, SplitLuc CETSA, lactate assay, biochemical assay and SPR. Compounds with no detected binding are annotated as "0", and compounds with no annotated data for a particular method are blank squares.* ***(B)*** *Thermal Dose-Response Curves for highlighted compounds from the $LDHA_i$ set. Curves shown are baseline-corrected data from single experiments.* ***(C)*** *Concentration-Response curves for highlighted compounds using RT-CETSA and analyzed by $T_{agg}$, AUC, and NPARC. Data shown are from n=3 biological replicates (mean ± SD).* ***(D)*** *Correlation between target engagement potency values calculated using SplitLuc CETSA or RT-CETSA (NPARC method), for twenty-nine LDHA inhibitors. Dotted line represents equipotency in the two assays.* ***(E)*** *Testing of the minimum significant ratio (MSR) and related parameters for the AUC metric. The mean ratio (MR) is shown as a solid blue line, Limits of Agreement (LsA) in dashed red lines, and ratio limits (RL) in dashed green lines.*

Supplementary Figure S4

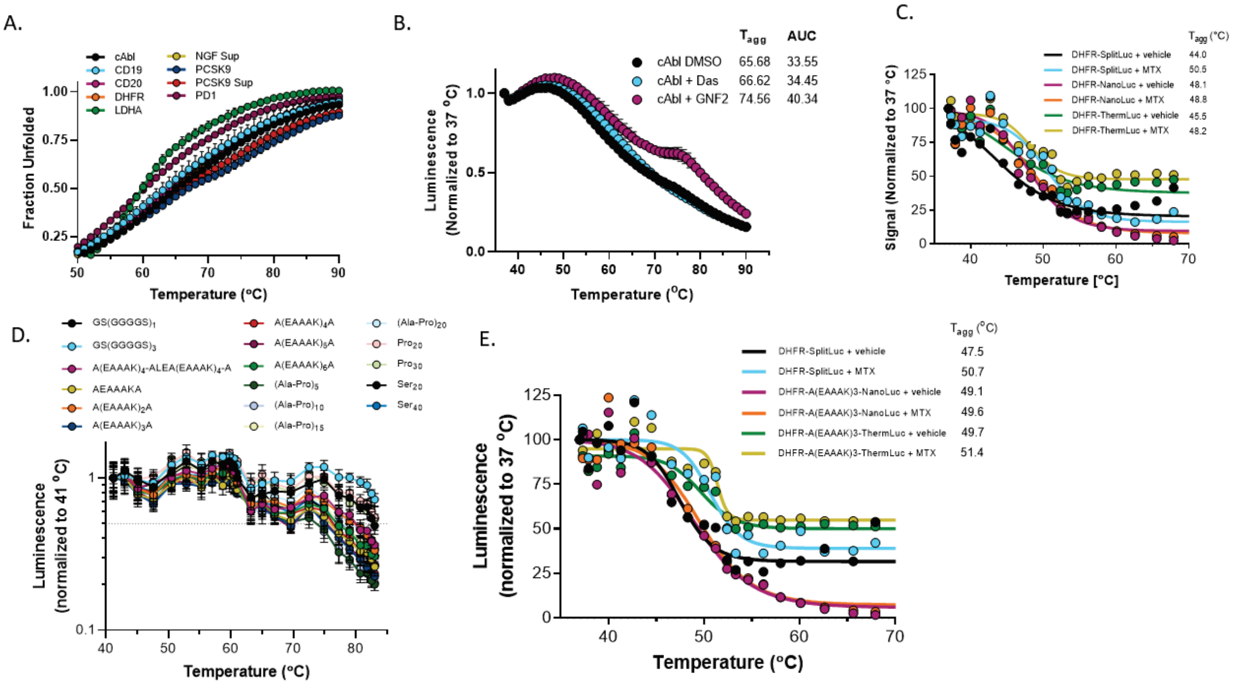


*Fig S4:* ***(A)*** *RT-CETSA analysis of various target-ThermLuc fusions (mean ± SD, n=32 wells). The secreted fraction (labeled 'sup') was examined for several proteins by transferring culture medium to a PCR plate before beginning the assay.* ***(B)*** *Target engagements of 10 µM Dasatinib (orthosteric inhibitor) or 40 µM GNF-2 (allosteric inhibitor) with cAbl-ThermLuc (ABL1, kinase-only domain) was examined by RT-CETSA (Mean ± SD, n=5).* ***(C)*** *Thermal melt profile of DHFR fusions (SplitLuc, NanoLuc, or ThermLuc), for vehicle or methotrexate (MTX) treated cells, examined using the endpoint CETSA method (mean, n=2).* ***(D)*** *Seventeen different linkers were inserted between DHFR and ThermLuc to examine effects on thermal stability in RT-CETSA (mean ± SD, n=3). Rank-ordering of melting temperatures is indicated in the legend.* ***(E)*** *Methotrexate target engagement was examined for DHFR fusions containing an A[EAAAK]$_3$ linker between the target and NanoLuc or ThermLuc (mean ± SD, n=2).*

Supplementary Movie 1: Monitoring thermal destabilization of LDHA-ThermLuc in HEK293T cells using real-time CETSA (RT-CETSA). Cells were treated with vehicle control or LDHA inhibitors, as indicated. Compound 63 (Rai et al.) labeling in the movie refers to compound **1** as numbered in this manuscript.
